# Structural neural connectivity analysis in zebrafish with restricted anterograde transneuronal viral labeling and quantitative brain mapping

**DOI:** 10.1101/764035

**Authors:** Manxiu Ma, Stanislav Kler, Y. Albert Pan

## Abstract

The unique combination of small size, translucency, and powerful genetic tools makes larval zebrafish a uniquely useful vertebrate system to investigate normal and pathological brain structure and function. While functional connectivity can now be assessed (via fluorescent calcium or voltage reporters) at the whole-brain scale, it remains challenging to systematically determine structural connections and identify connectivity changes during development or disease. To address this, we developed <u>T</u>racer with <u>R</u>estricted <u>A</u>nterograde <u>S</u>pread (TRAS), a novel vesicular stomatitis virus (VSV)-based neural circuit labeling approach. TRAS makes use of replication-incompetent VSV (VSVΔG) and a helper virus (lentivirus) to enable anterograde transneuronal spread between efferent axons and their direct postsynaptic targets but restricts further spread to downstream areas. We integrated TRAS with the Z-Brain zebrafish 3D atlas for quantitative connectivity analysis and identified targets of the retinal and habenular efferent projections, in patterns consistent with previous reports. We compared retinofugal connectivity patterns between wild-type and *down syndrome cell adhesion molecule-like 1* (*dscaml1*) mutant zebrafish and revealed differences in topographical distribution and potential changes in the retinofugal targeting of excitatory versus inhibitory retinorecipient cells. These results demonstrate the utility of TRAS for quantitative structural connectivity analysis that would be valuable for detecting novel efferent targets and mapping connectivity changes underlying neurological or behavioral deficits.

## Introduction

The function of the brain is closely linked to its structure, how its billions of constituent cells are wired and connected by trillions of synapses. Understanding how these connections are formed and maintained is key to gaining mechanistic insight towards brain function and identifying the causes and treatments for neuropsychiatric disorders (Belmonte et al., 2004; Lynall et al., 2010; Fornito et al., 2015). Techniques for mapping the structure and function of the brain have progressed rapidly in past decades, from anatomical structural analysis to functional computation, and from human participants to animal models (Kasthuri et al., 2015; Glasser et al., 2016). However, the large number of neurons and vast spatial scale of neuronal structures (from meters to nanometers) of the mammalian brain makes mapping neuronal networks at the cellular level and correlating them with development and disease a daunting task (Swanson and Lichtman, 2016).

Zebrafish (*Danio rerio*), a small tropical fish, has emerged as an accessible model for studying behavior, neuronal networks, and cellular connectivity (Orger and de Polavieja, 2017). Zebrafish has analogous neuroanatomy and neurochemistry to mammals and can perform complex sensory, motor, and cognitive functions early during larval stages (5-10 days post fertilization, dpf). Importantly, at this stage, there are only roughly 100,000 neurons in the brain, 80% of which can be imaged and physiologically recorded in live, behaving animals (Ahrens et al., 2013; Chen et al., 2018; Abdelfattah et al., 2019). As a result, zebrafish whole-brain functional imaging studies have been able to generate cellular resolution neuronal activity maps under different behavioral contexts and linking activity maps to disease states such as autism spectrum disorder and epilepsy (Sakai et al., 2018; Thyme et al., 2019). The ability to fully interpret patterns of functional connectivity and determine causality for disorders, however, is limited by the lack of detailed structural information on neuronal wiring in zebrafish, which still lags behind other commonly used model organisms like mice, *Drosophila*, and *C. elegans*. Efforts are ongoing to map the full complement of neuronal connections (i.e., connectome) with electron microscopy in the larval and adult zebrafish, but the time and labor-intensive nature of synapse-level reconstruction has thus far restricted investigations to connections within smaller brain regions (Wanner et al., 2016; Hildebrand et al., 2017; Vishwanathan et al., 2017; Svara et al., 2018).

To address this, we sought to develop a virus and light imaging-based structural mapping technique that would allow for quantitative brain-wide mapping of neuronal connectivity in larval zebrafish. Previously, we found that recombinant vesicular stomatitis virus (VSV) can function as an anterograde transsynaptic tracer in a wide range of organisms (Mundell et al., 2015). VSV is a negative-strand RNA virus in the *Rhabdoviridae* family, which also includes the rabies virus (RABV). In contrast to RABV, which spreads retrogradely (from dendrite to presynaptic afferent axons), VSV spreads anterogradely (from efferent axons to their postsynaptic targets) when enveloped by its endogenous glycoprotein (VSV-G) (Beier et al., 2011b; Mundell et al., 2015). VSV injection into the retina of mice, chicken, and larval zebrafish lead to highly efficient labeling of the optic nerve and targets of the visual pathway, including both direct retinorecipient connections and areas further downstream. These findings open the door for utilizing VSV for zebrafish structural circuit mapping. Substantial limitations, however, still remain. First, the spread of VSV is unrestricted, making it difficult to disambiguate direct (monosynaptic) and indirect (polysynaptic) connections. Second, replication-competent VSV and VSV-G expression are cytotoxic and lead to rapid deterioration of health in larval zebrafish (Hoffmann et al., 2010; Mundell et al., 2015). Finally, there is no established method for quantifying and annotating viral labeling in zebrafish, which is necessary to correlate anatomical tracing with functional imaging.

In this study, we developed a novel approach utilizing replication-incompetent VSV to achieve restricted anterograde transneuronal spread in zebrafish. We also developed an imaging and processing pipeline to register 3D image stacks to the widely used and extensively annotated Z-Brain digital atlas (Randlett et al., 2015). This method, termed TRAS (<u>T</u>racer with <u>R</u>estricted <u>A</u>nterograde <u>S</u>pread), allows for a quantitative description of efferent connectivity based on neurotransmitter types and specific locations. We applied TRAS to investigate the axon projection patterns of retinorecipient cells and identified potential connectivity changes in zebrafish carrying a mutation in *down syndrome cell adhesion molecule-like 1 (dscaml1)*, a causal gene for autism spectrum disorder and human cortical abnormalities (Fuerst et al., 2009; Iossifov et al., 2014; Karaca et al., 2015; Galicia et al., 2018; Ma et al., 2019).

## Materials and Methods

### Zebrafish husbandry

Zebrafish (all ages) were raised under a 14/10 light/dark cycle at 28.5°C. Embryos and larvae were raised in water containing 0.1% Methylene Blue hydrate (Sigma-Aldrich). With the exception of *nacre* mutants, embryos were transferred to E3 buffer containing 0.003% 1-phenyl-2-thiourea (PTU; Sigma-Aldrich) to prevent pigment formation at 24 hours post-fertilization. Developmental stages are as described by Kimmel et al. (Kimmel et al., 1995). Sex was not considered as a relevant variable for this study, as laboratory zebrafish remain sexually undifferentiated until two weeks of age, beyond the stages being used (0-9 dpf) (Maack and Segner, 2003; Wilson et al., 2014). All experimental procedures are performed in accordance with Institutional Animal Care and Use Committee guidelines at Augusta University and Virginia Tech.

### Transgenic and mutant zebrafish lines

The *dscaml1^vt1^* mutant line was generated by TALEN-targeted mutagenesis, which resulted in a seven base pair deletion and subsequent early translational termination (Ma et al., 2019). The *Tg(elavl3:H2B-GCaMP6f)* line was generously provided by E. Aksay at Weill Medical College, with permission from M. Ahrens at HHMI Janelia Farm Research Campus (Kawashima et al., 2016). The *vglut2a:GFP* line [*Tg(slc17a6b:EGFP)*] was generously provided by J. Fetcho at Cornell University with permission from S. Higashijima at the National Institute for Basic Biology (Bae et al., 2009).

### Preparation of VSVΔG

VSV was prepared using methods detailed by Beier et al. 2016 (Beier et al., 2016). 293T cells (ATCC, #CRL-3216) were transfected at 80% confluency on 75 cm^2^ flasks with 7 μg of *pCI-VSVG* plasmid (Addgene, #1733) and incubated overnight at 37°C. Afterward, cells were infected with VSVΔG-RFP (VSVΔG for short) (Beier et al., 2011b) at a multiplicity of infection (m.o.i.) of 0.1. Viral supernatants were collected for the subsequent three days at 24-hour intervals and combined. Cell debris was precipitated by centrifugation at 1,000 g for 20 min. To concentrate VSVΔG, viral supernatant was ultracentrifuged for three hours at 80,000 g with an SW32Ti rotor, and the pellet was resuspended in 100 μl of culture medium. Viral stocks were titered by serial dilution on 90% confluent 293T cells. The number of fluorescent foci was calculated at two days post infection (dpi) by identifying RFP-positive cells. Typical viral titer was higher than 1×10^9^ focus forming units/ml (ffu/ml).

For *in vitro* trans-complementation of VSVΔG with lentivirus, BHK-21 cells (ATCC, #CCL-10) were seeded into 96-well plate and incubated overnight to reach 20,000 cells per well. Cell culture was co-infected with VSVΔG (m.o.i.=0.005) and VSV-G pseudotyped lentivirus (m.o.i.=0-10,000) (lentivirus-SIN-CMV-eGFP or lentivirus-SIN-Ubi-iCre-mCherry; GT3 Core Facility of the Salk Institute). At two hours post infection, cells were washed twice with PBS and incubated with fresh medium supplemented with 2% serum. At 2 dpi, the spread of VSV was visualized by fluorescent microscopy. The media in each well were collected and titered to evaluate viral yield.

### Virus injection

Viral injections were performed as previously described (Mundell et al., 2015; Beier et al., 2016). Briefly, glass capillaries (TW100F-4; World Precision Instruments) were pulled into injection needles with a pipette puller (P-97; Sutter Instruments). The tips of injection needles were trimmed to create a ~10 μm opening. Virus injection solution was made by diluting VSVΔG and lentivirus stock with tissue culture medium (DMEM; Fisher Scientific), with Fast Green dye (BP123-10; Fisher Scientific) as the injection marker. 2 μl of injection solution was loaded into the injection needle with a Microloader pipette tip (930001007; Eppendorf), and mounted into a microelectrode holder connected to a pneumatic PicoPump (PV820; World Precision Instruments). Injection volume was determined by calibrating the volume of the injected droplet on a stage micrometer (50-753-2911; Fisher Scientific). The hold pressure of the PicoPump was adjusted so that there was a slight outflow of virus solution when the needle tip was immersed in fish water.

For retina injection, 2.5 or 3 dpf larvae were anesthetized in Tricaine (0.013% w/v, AC118000500; Fisher Scientific) mounted laterally inside the center chamber of a glass-bottom dish (P50G-1.5-14-F; MatTek) with 1.5% low melting-point agarose (BP1360; Fisher Scientific). After the agarose has solidified, the dish is filled with Tricaine solution (0.013%) to maintain anesthesia. Under a stereo dissecting microscope (SMZ18; Nikon), the needle tip was moved with a micromanipulator (MN-151; Narishige) to approach the fish from the rear and penetrated the temporal retina, with the needle tip being in the neural retina. 0.25-0.5 nl of virus solution (concentrations as described in the Results section) were injected inside the retina. After injection, larvae were recovered from the agarose and returned to a 28°C incubator. Overall, the combination of VSVΔG and lentivirus (i.e., TRAS) only labeled cells in areas innervated by or adjacent to the optic nerve. It is worth noting that in one experiment we did observe neuronal cell bodies in the hindbrain and contralateral midbrain, which are two synapses downstream from RGCs (Helmbrecht et al., 2018) (1-2 cells in 50% of injected fish, n=10, Supplementary Figure S1). This may be due to VSVΔG self complementation in the retinorecipient cells at high m.o.i. For all quantitative analysis involving selective labeling for primary retinorecipient cells, we used viral titers that did not result in secondary spread (VSVΔG at 3×10^7^ ffu/ml and lentivirus at 3×10^10^ ffu/ml).

For habenula injection, 3 dpf larvae were mounted as described for retina injection, with the dorsal side up. The agarose and skin above the left habenula were carefully removed with a sharpened tungsten needle (10130-05; Fine Science Tools). The injection needle tip was inserted into the left habenula and remained there for 5 s, allowing the slow outflow of virus solution (VSVΔG at 3×10^8^ ffu/ml, lentivirus at 1×10^11^ ffu/ml) to immerse the surrounding tissue. At 1 dpi, larvae with habenula-specific RFP expression were screened and later fixed at 3 dpi for immunohistochemistry and confocal imaging.

### Immunohistochemistry

Whole-mount immunohistochemistry was performed as described by Randlett et al. 2015 (Randlett et al., 2015). Zebrafish larvae were fixed overnight with 4% PFA and 0. 25% Triton X-100 (Fisher Scientific) in 1X PBS (diluted from 10%PFA; Polysciences), then washed with 1X PBS and 0. 25% Triton X-100. H2B-GCaM6f was stained with FluoTag-X4 anti-GFP (N0304-At488; NanoTag); GABA was stained with Rabbit anti-GABA (A2052; Sigma-Aldrich); ERK1/2 was stained with mouse anti-ERK1/2 (4696S; Cell Signaling Technology). The sRIMS solution, which is D-Sorbitol (Sigma-Aldrich) dissolved in PBS with 0.1% Tween-20 (Fisher Scientific) and 0.01% Sodium Azide, was used for optical clearing (Yang et al., 2014). Samples are immersed for 15 min (or until sunken to the bottom of the Eppendorf tube) through a gradient series (8.75% to 70%) of D-Sorbitol.

### Image acquisition

Epifluorescence images of cultured cells were acquired under a Nikon Eclipse Ts2 inverted fluorescent microscope. Images of zebrafish were acquired using a Nikon A1 laser scanning confocal system with a CFI75 Apochromat LWD 25x water-immersion objective. For TRAS quantification, image stacks were acquired at a standard resolution of 0.49 x 0.49 x 2.0 μm^3^ per voxel. For efferent tract tracing, a standard resolution stack and a high-resolution stack (0.38 x 0.38 x 0.5 μm^3^ per voxel) were acquired for each fish.

### Efferent tract tracing

Standard resolution image stacks were morphed to the Z-Brain *elavl3:H2B-RFP* template using CMTK (Randlett et al., 2015). High-resolution stacks were then morphed to its corresponding low-resolution stacks to register to Z-Brain coordinates. Morphed high-resolution stacks were imported into the neuTube software for tracing, in accordance with the neuTube online manual (https://www.neutracing.com/manual/) (Feng et al., 2015). The SWC files were saved and imported into Fiji plugin “Simple Neurite Tracer” then saved as an overlay.

### TRAS quantitation with Z-Brain

Image stacks used for cell-type characterization were morphed to the Z-Brain *elavl3:H2B-RFP* template using CMTK, using the GCaMP6f channel as reference (Randlett et al., 2015). After morphing, the Fiji software’s ROI manager (Analyze>Tools>ROI manager; extra-large size dot) was used on the VSVΔG channel to mark all VSVΔG+ cells. The marked positions (ROIs) were saved into a zip file and overlaid onto the GCaMP6f channel. The ROIs that were GCaMP6f-negative were removed so that the remaining ROIs represented TRAS-labeled neurons (neuronal ROIs). Next, the neuronal ROIs were overlayed onto the GABA channel to create two subsets: the GABA+ ROIs (inhibitory neurons) and GABA-ROIs (excitatory neurons, created by subtracting the neuronal ROI with the GABA+ ROIs). Lastly, these two ROIs were overlaid onto a Z-brain reference-sized (X:Y:Z=1121×496×276 μm^3^ at 0.8×0.8×2 μm^3^ per voxel) blank stack, with inhibitory neuron ROIs pseudocolored magenta and excitatory neuron ROIs pseudocolored green.

To quantitate the anatomical distribution of retinorecipient cells, we followed the procedures for Z-Brain MAP-map analysis, but with several modifications (Randlett et al., 2015). Instead of using the MakeTheMAPmap.m MATLAB script, we used a custom Fiji macro script to create an image file. The image file which was quantified using a modified ZBrainAnalysisOfMAPmaps.m script that quantify sum pixel intensity values for neuronal ROIs within each region mask. The output Intensity values were converted to cell counts, based on an estimate of the pixel values generated from a single-cell ROI (~18,265). For heatmap display and cohort-wise comparison of individual regions, the intensity signals for each fish were normalized by the total signal from “Diencephalon” and “Mesencephalon.” The numbers were then imported into MATLAB to make a scaled color map using “imagesc” function.

### Analysis of topographical distribution

To analyze the topographical distribution of wild-type and *dscaml1−/−* retinorecipient cells (Fig. 5), the ROI files from all fish within a cohort were combined into a single zip file and overlayed into a Z-Brain compatible blank stack, as described previously for single fish ROI image files. ROI files from individual fish from the same genotype were combined into a single zip file. The x-y-z coordinates of each ROI dot were used in “scatter3” function in MATLAB to create 3-D scatter plots. The same coordinates were imported into GraphPad for analysis of distribution properties. The cumulative frequency statistics were done using the K-S test provided within the Prism software (GraphPad).

### Excitatory/Inhibitory ratio

The total number of neurons in the *dscaml1−/−* cohort (267) was normalized to the total number of wild-type neurons (340), and the normalized increase of 27.34% in mutants was proportionally applied to individual anatomical regions and rounded up to a whole number. Excitatory-to-inhibitory neuron number ratios were calculated based on the estimated number of neurons within each anatomical region. In order to obtain valid ratio values, only regions with non-zero values were used for ratio analysis.

## Results

### Trans-complementation of VSVΔG by VSV-G coated lentivirus

For both VSV and RABV, the envelope glycoprotein (G) gene is essential for binding, internalization, membrane fusion, and release of the viral genome into the host cell (Albertini et al., 2012; Kim et al., 2017). A recombinant virus with genomic deletion of the G gene (ΔG) can infect and replicate inside the cell but is unable to spread, unless the host cell complements the virus by providing G *in trans* (Wickersham et al., 2007; Beier et al., 2011b). Trans-complementation can, therefore, be utilized to restrict viral spread to direct synaptic partners. For instance, expressing the RABV-derived glycoprotein (RABV-G) in neurons at the injection site (starter cells) allowed VSVΔG or RABVΔG to spread from the starter cells to their input neurons. Once inside the input neurons, the virus can no longer spread (Wickersham et al., 2007; Beier et al., 2013).

Given VSV’s ability to spread anterogradely across synapses, we asked whether trans-complementing VSVΔG virus with VSV glycoprotein (VSV-G) could enable restricted anterograde spread (Mundell et al., 2015). Our efforts to express VSV-G *in vivo* through transgenesis was unsuccessful, possibly due to the pathogenic effects of VSV-G when persistently expressed (Yee et al., 1994). As an alternative, we tested whether VSV-G protein could be transduced to cells directly. We took advantage of the fact that most commercially available lentiviruses are coated with VSV-G and examined *in vitro* whether concomitant VSVΔG/lentivirus infection could provide sufficient VSV-G to trans-complement VSVΔG. 293T cells were co-infected with VSV-G enveloped, RFP-expressing VSVΔG at low density [multiplicity of infection (m.o.i.) = 0.005] and lentivirus at a range of densities (m.o.i. = 0 to 1,000). At two days post-infection (dpi), we visualized the spread of VSVΔG by fluorescent microscopy and determined the yield of newly synthesized VSVΔG in the media. Indeed, lentivirus complemented VSVΔG in a dose-dependent manner, indicating that VSV-G on the envelope of lentivirus could be taken up by VSVΔG to form functional virions (Fig. 1A-C).

**Figure. 1.**
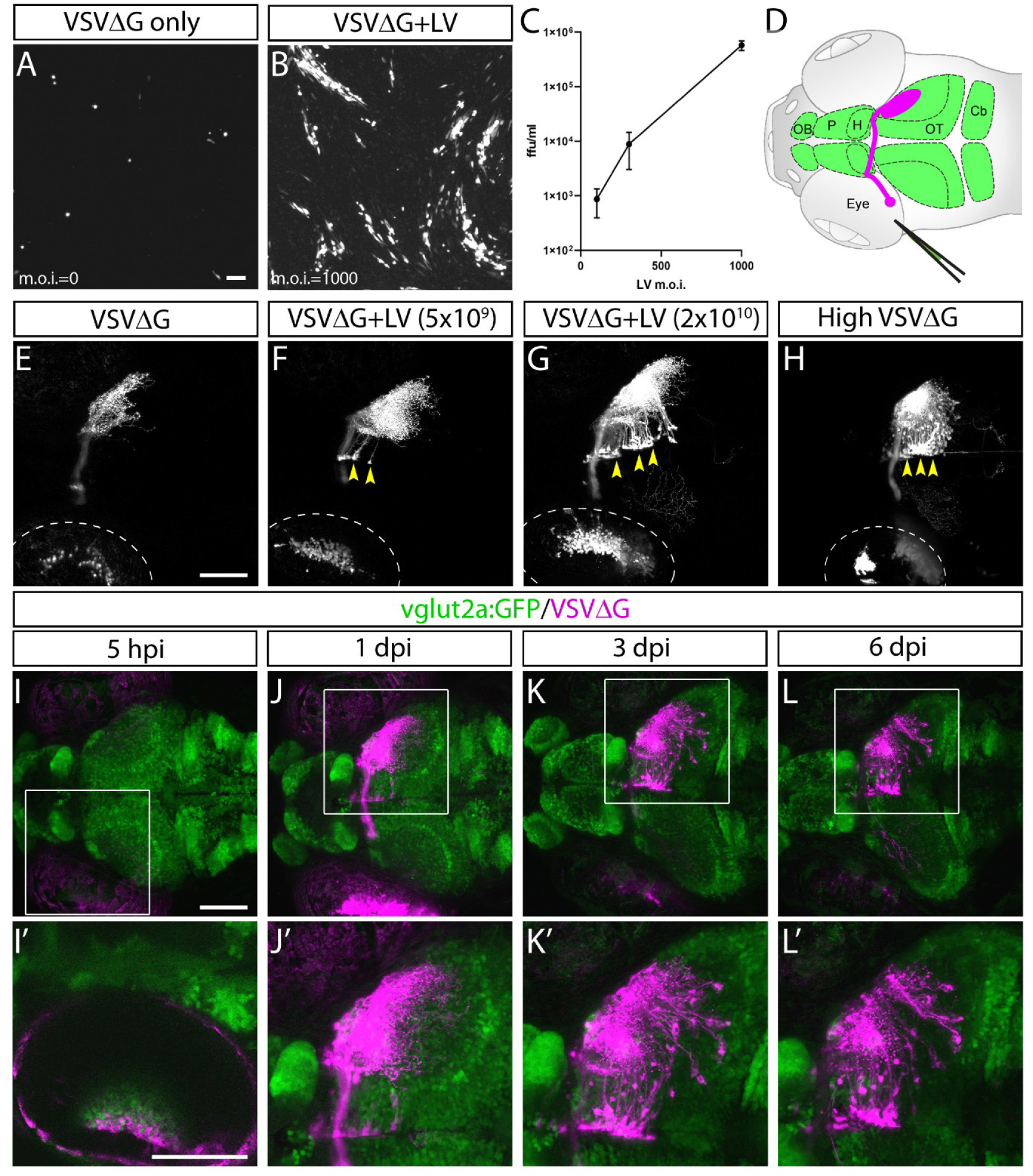
Lentivirus enabled in vitro and in vivo trans-complementation of VSVΔG and transneuronal spread. **A-C**, lentivirus trans-complementation in vitro. VSVΔG was able to infect 293T cells but was unable to spread to neighboring cells, as evident by sparse single-cell infections. (A). In conjunction with lentivirus, VSVΔG was able to both infect and spread, as evident by the presence of large infected plaques (B). The extent of VSVΔG amplification (as measured by viral titer) is positively correlated with lentivirus titer, expressed in m.o.i. (C). **D,** Illustration of viral injection and labeling of the optic nerve. Virus was microinjected into the left eye, which infected the RGC and resulted in fluorescent labeling of the RGC axons (i.e., optic nerve, magenta). The layout of the larval CNS is labeled in green, with the olfactory bulb (OB), pallium (P), habenula (H), optic tectum (OT), and cerebellum (Cb) labeled. **E-H,** In vivo trans-complementation and spread of VSVΔG by lentivirus. In the absence of lentivirus, VSVΔG infected RGCs and fluorescently labeled the optic nerve, but no spread in the CNS was observed (E). When lentivirus and VSVΔG were coinjected, cellular labeling was observed in the CNS (yellow arrowheads), indicating transneuronal spread. Similar patterns of spread was also seen at very high VSVΔG titer (2×109 versus 108 ffu/ml for panels E-G), suggesting that VSVΔG was able to self-complement (H). **I-L,** Time course of VSVΔG infection and spread with lentivirus trans-complementation, with RFP expression from VSVΔG (magenta) and GFP expression from the *vglut2a:GFP* transgene (green). Box regions are shown at higher magnifications below (I’-L’). Scale bars are 100 μm. Images in the same row are shown at the same scale.

Lentivirus-mediated trans-complementation was also effective *in vivo*, enabling transneuronal spread. By itself, VSVΔG injection (0.5 nl at 10^8^ ffu/ml) into the eye resulted in retina infection and RFP labeling of the optic nerve, but no cellular labeling in the brain (Fig. 1D-E). This suggested that VSVΔG was not released from axon terminals to initiate a new cycle of infection in the brain. When low (5×10^9^ ffu) or high titer lentivirus (2×10^10^ ffu) was co-injected with VSVΔG (10^8^ ffu/ml), we observed cellular labeling in the brain in both conditions, with more spread in injections with high titer lentivirus (Fig. 1F-G). This agrees with our *in vitro* results and indicates that high titer lentivirus can trans-complement VSVΔG, allowing viral spread from axon terminals.

The ability of lentivirus to trans-complement was not dependent on what the lentivirus genome encodes. Two types of VSV-G coated lentivirus were tested (lentivirus-SIN-CMV-eGFP and lentivirus-SIN-Ubi-iCre-mCherry). Both were able to mediate trans-complementation, and neither were able to drive transgene expression (eGFP and iCre-mCherry, respectively) on their own in fish at 6 dpi. Lastly, we asked whether VSVΔG could self-complement at higher titer since VSVΔG itself was also enveloped in VSV-G. Indeed, high titer VSVΔG (2×10^9^ ffu/ml) could spread from the RGC to retinorecipient cells in the brain (Fig. 1H). Together, these results show that VSV-G from different viral particles could be recycled to form infectious VSV particles.

### Restricted anterograde spread of VSVΔG in the zebrafish visual pathway

Since lentivirus was supplied at the injected site, only neurons at the injection site (starter cells) should be able to mediate spread. The spread from the starter cells should be limited to direct postsynaptic targets, i.e., anterograde monosynaptic spread. To test this, we examined whether the spatial and temporal patterns of VSVΔG spread were consistent with monosynaptic spread from retinal ganglion cells (RGCs) to retinorecipient neurons in the brain.

VSVΔG and lentivirus were coinjected into anesthetized 3 dpf zebrafish larvae, followed by live confocal imaging at different time points (n=8 animals). Initial RFP expression from VSVΔG was present in the injected (left) eye as early as 5-hours post infection (hpi) (Fig. 1I). Cellular labeling in the contralateral (right) brain was observed at 1 dpi, and more cells were labeled at 3 dpi (Fig. 1J-K). At 6 dpi, there was no further spread to other brain regions, compared to 3 dpi (Fig. 1L). This pattern of labeling is distinct from non-G-deleted (replication competent) VSV, which rapidly progressed from axonal labeling to cell body labeling in downstream areas like the cerebellum and habenula at 3 dpi (Mundell et al., 2015). These results suggest that lentivirus trans-complementation primarily mediated anterograde monosynaptic spread. We call this new technique <u>T</u>racer with <u>R</u>estricted <u>A</u>nterograde <u>S</u>pread (TRAS, pronounced like *trace*).

### Efferent projections of retinorecipient cells were revealed by TRAS

Retinorecipient cells extend axons to different parts of the brain to mediate visually guided cognitive, sensory, motor, and homeostatic functions. Previous studies have utilized transgenic reporter lines to characterize efferent projections of subsets of retinorecipient cells, but there has not been a method that could unbiasedly label retinorecipient cells in different brain regions and reveal their efferent projections (Zhang et al., 2017; Helmbrecht et al., 2018; Kramer et al., 2019). With TRAS, we observed several prominent efferent tracts from retinorecipient neurons, innervating the telencephalon (6 of 8 animals), habenula (2 of 8), midbrain tegmentum (7 of 8), contralateral optic tectum (7 of 8), cerebellum (6 of 8), and along the ventral hindbrain (6 of 8) (Fig. 2, image stack for panel 2A is shown in supplementary video 1). These projection patterns are reminiscent of the efferent projections of tectal/pretectal retinorecipient neurons, further supporting the idea that TRAS primarily labels retinorecipient cells (Sato et al., 2007; Mundell et al., 2015; Helmbrecht et al., 2018; Kramer et al., 2019). The projection into the telencephalon by retinorecipient neurons, to our best knowledge, has not been reported previously. These axon projections extend rostrally to enter the subpallium and then courses dorsally to the caudal pallium. Some axons crossed near the anterior commissure. These pallium-projecting neurons may serve similar roles as the mammalian lateral geniculate neurons to relay sensory information to higher visual areas (Mueller, 2012). These results show that TRAS could be used to identify not only postsynaptic cells but also downstream areas innervated by these cells.

**Figure 2.**
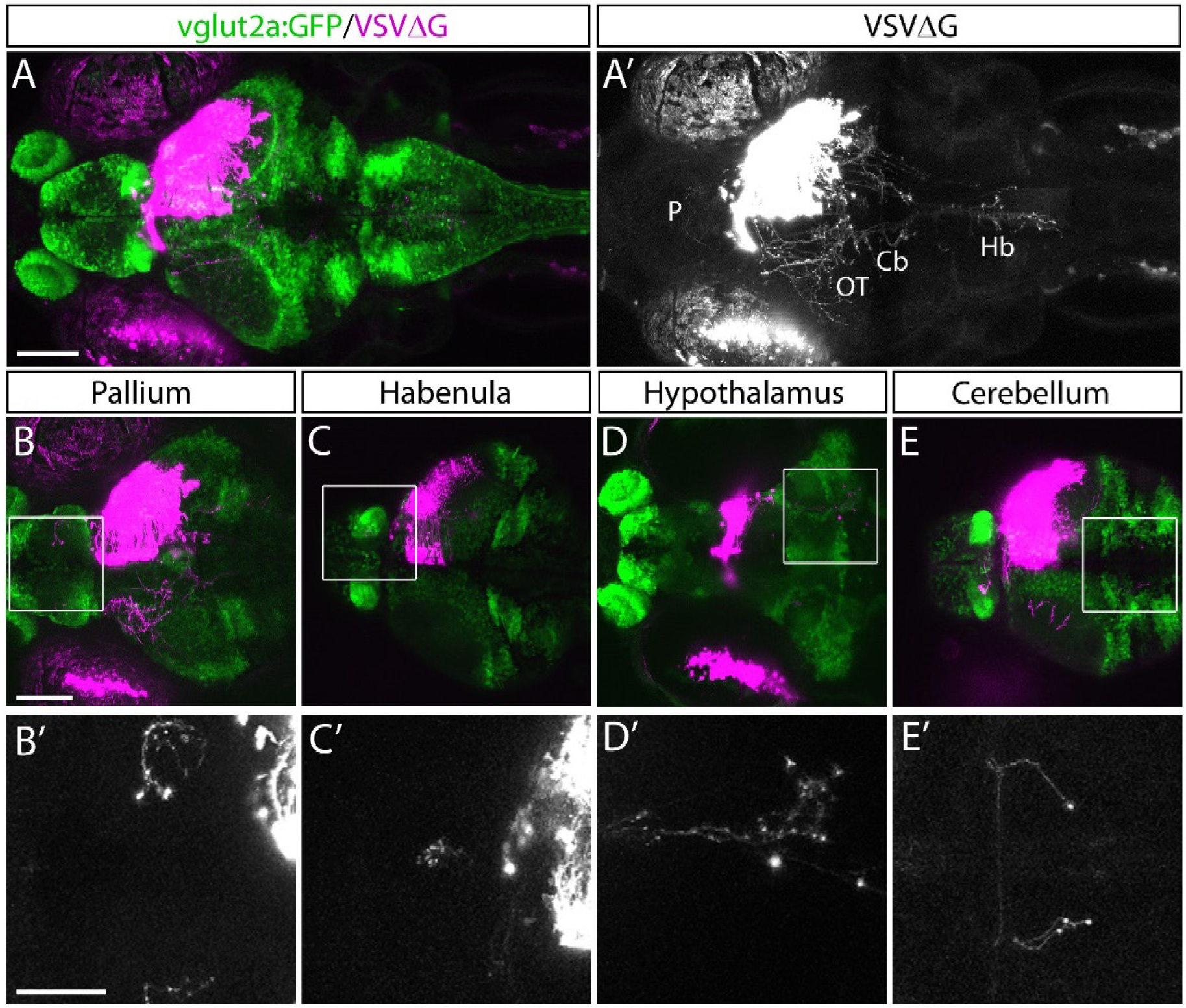
Efferent projections of retinorecipient cells. **A-A’,** Confocal maximal intensity projection (dorsal view) of TRAS-labeled larva, with RFP expression from VSVΔG in magenta (A) or white (A’). Axon projections can be seen in the pallium (P), optic tectum (OT), cerebellum (Cb), and hindbrain (Hb). **B-E,** Maximal intensity projection confocal substacks that contained the pallium (B), habenula (C), hypothalamus (D), and cerebellum (E). RFP expression from VSVΔG (magenta) and GFP expression from the *vglut2a:GFP* transgene (green) are shown in B-E, while boxed region is shown at higher magnification in B’-E’, with only the RFP channel (white). Scale bars are 100 μm in A and B, and 50 μm in B’. Images in the same row are shown at the same scale.

### 3D mapping and cell-type characterization

To quantify connectivity patterns, we registered TRAS-labelled image stacks to the Z-Brain zebrafish brain atlas (Randlett et al., 2015). Transgenic fish expressing neuronal-localized nuclear GCaMP6f (*elavl3:H2B-GCaMP6f*) were injected at 2.5 dpf, into the temporal region of the left eye (representing the frontal visual field). Infected larvae were fixed at 3 dpi and stained with anti-GFP (to amplify the GCaMP6f signal) and anti-GABA. Stained samples were then cleared in sRIMS, a sorbitol-based mounting media that was crucial to resolving single cells and axon tracts in the ventral brain (Yang et al., 2014) (Fig. 3A-B, Supplementary Figure S2). By extracting the Z-Brain *elavl3:H2B-RFP* stack from ZBrainViewer as the reference, image stacks were morphed and aligned with the Computational Morphometry Toolkit (CMTK) (Rohlfing and Maurer, 2003; Jefferis et al., 2007; Randlett et al., 2015).

**Figure 3.**
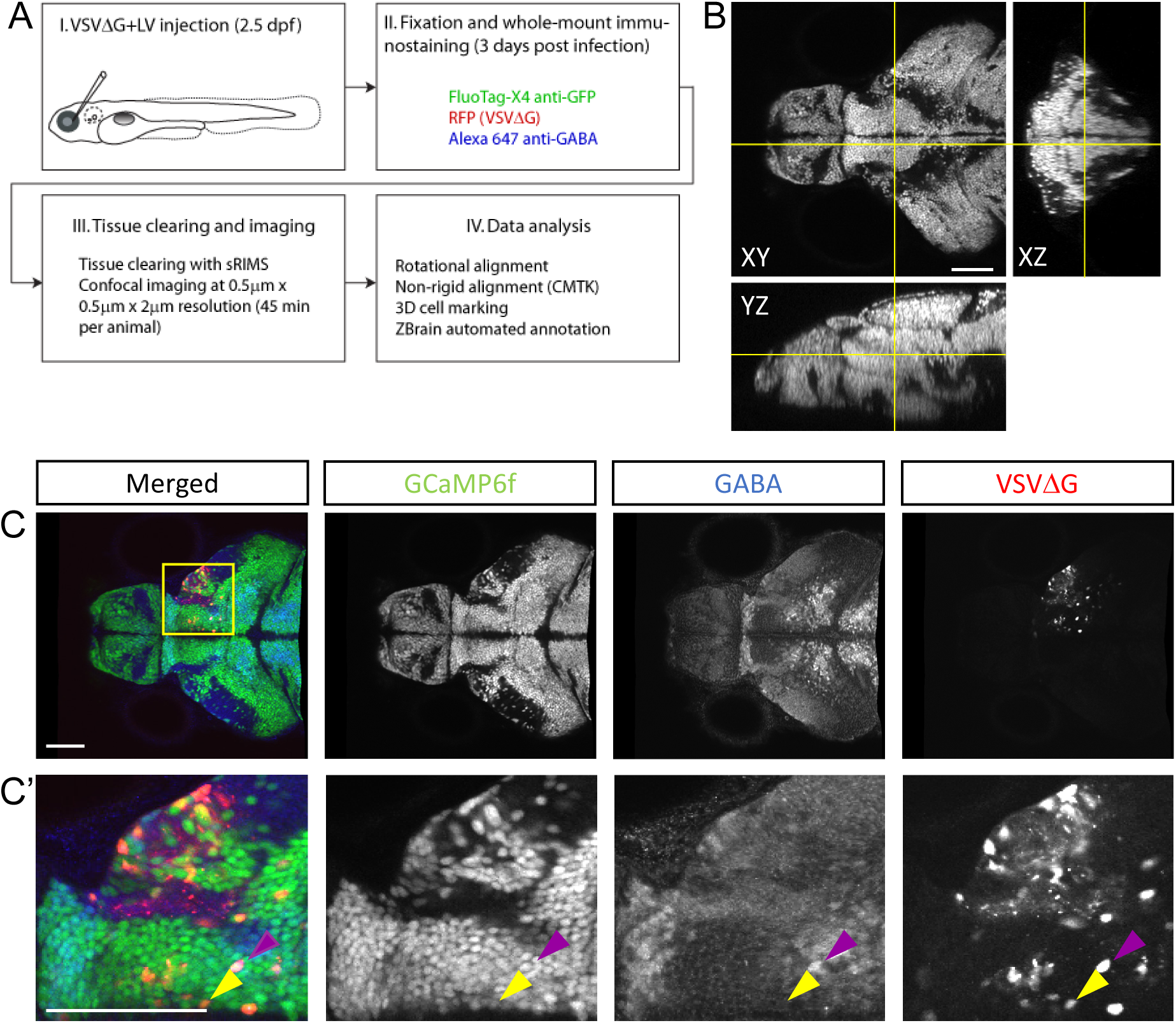
Cell-type characterization of TRAS labeling. **A,** The workflow for TRAS labeling, tissue processing, image acquisition, and data analysis. **B,** Orthogonal views of an imaged fish after tissue clearing with sRIMS. Orthogonal views (XY, XZ, and YZ) of confocal image stacks are shown. **C-C’,** Cytochemical characterization of TRAS-labeled cells. A single confocal imaging plane is shown, with merged, GCaMPf, GABA, and VSVΔG channels as indicated. Boxed area in C is shown in higher magnification in C’. The purple arrowhead marks a GCaMP6f+/GABA+/VSVΔG+ inhibitory neuron. The yellow arrowhead marks a GCaMP6f+/GABA-/VSVΔG+ excitatory neuron. Scale bars are 100 μm.

To distinguish between different cell types, both transgenic and immunohistochemical markers were used. Nuclear GCaMP6f (expressed only in neurons) was used to distinguish between neuronal (GCaMP6f+) and non-GCaMP (GCaMP6f-, non-neuronal cells and HuC-neurons) cells. Anti-GABA staining was used to distinguish between non-GABAergic and GABAergic (inhibitory) neurons (Cui et al., 2005) (Fig. 3C). Since glycinergic neurons are not present in the retinorecipient areas, non-GABAergic retinorecipient neurons are predominantly excitatory. After CMTK, all TRAS labeled cells were converted into Z-Brain coordinates and categorized into three types: excitatory neurons (GCaMP6f+, GABA-) inhibitory neurons (GCaMP6f+, GABA+), and non-GCaMP cells. In total, 24 wild-type and 26 *dscaml1−/−* fish (see next section) were analyzed (Fig. 4A-B, supplementary video 2, supplemental Figure S3). The overall ratio of excitatory versus inhibitory cells was similar between wild-type and *dscaml1−/−* cohorts (p=0.515, Chi-square= 0.424). The annotated stack could be viewed with the Z-Brain off-line viewer and overlaid with anatomical annotations and reference stacks therein (Randlett et al., 2015).

**Figure 4.**
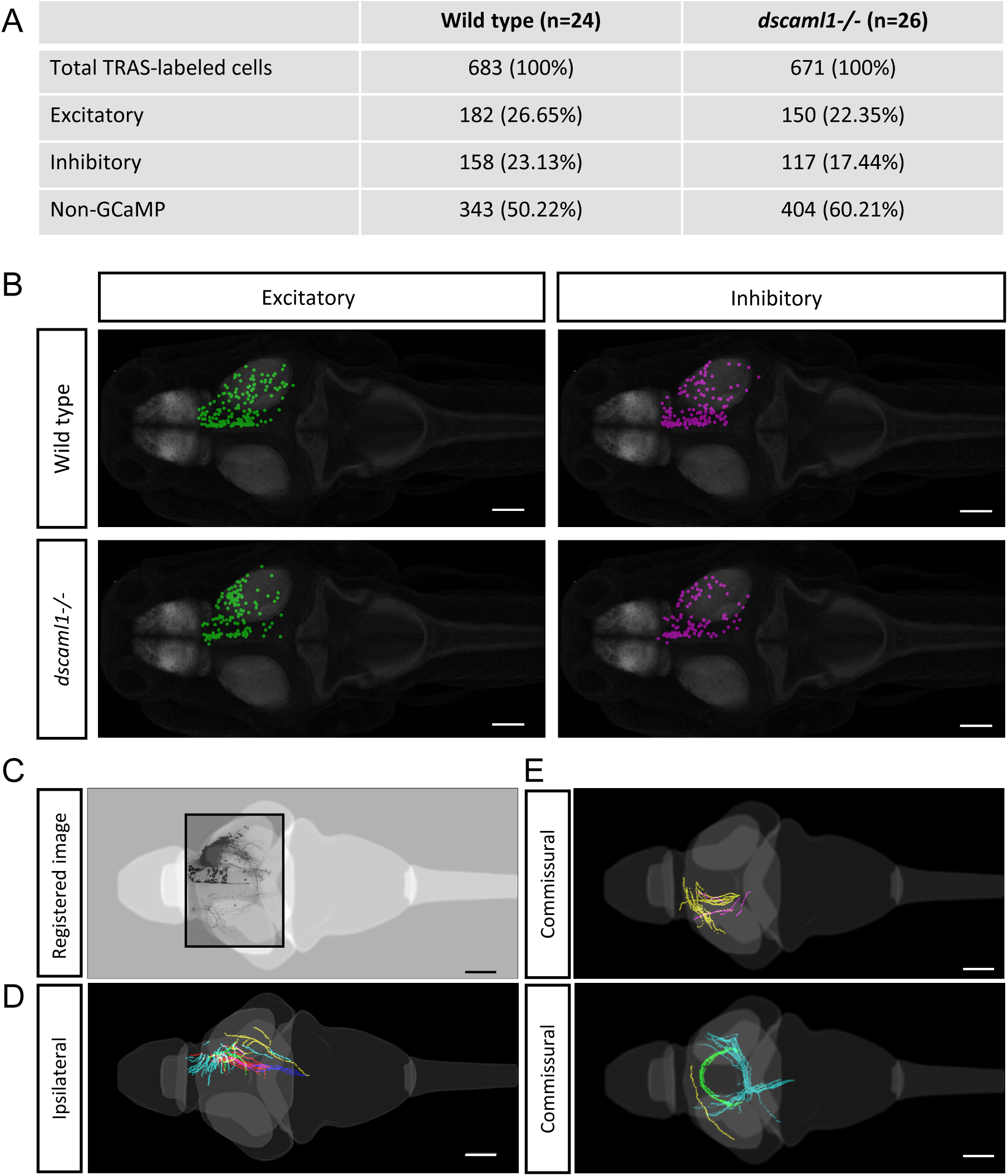
Annotation of TRAS-labeled neurons and efferent axons in the Z-Brain standard brain reference. **A,** Overview of all annotated TRAS-labeled retinorecipient cells within the wild-type and *dscaml1−/−* cohorts. **B,** Spatial layout of TRAS-labeled neurons (dorsal view, rostral to the left) overlayed onto the Z-brain reference brain scale, for wild-type (top row) and *dscaml1−/−* (bottom row) cohorts. Green dots mark excitatory neurons, magenta dots mark inhibitory neurons. **C-E,** Efferent tract tracing from wild-type larvae (n=10). Maximum Z-projection is shown for confocal image (C), traced ipsilateral tracts (D), and commissural tracts (E). Axons with similar trajectories are displayed in the same color. Scale bars are 100um.

We also verified that Z-Brain registered stacks could be used as templates for tracing the efferent projections of retinorecipient cells (Fig. 4C). We acquired both high-resolution and low-resolution image stacks for the same fish and used the high-resolution image stacks for tracing and low-resolution image stacks as the template to register to Z-Brain. We observed ipsilateral and commissural axon tracts, with morphologies that are similar to the tectal efferent tracts described in previous studies (Sato et al., 2007; Helmbrecht et al., 2018).

### Comparative analysis of retinofugal connectivity

In addition to normal patterns of retinofugal connectivity, TRAS and Z-Brain can be used to investigate retinofugal connectivity patterns in mutants with visual deficits. We focused on *Down Syndrome Cell Adhesion Molecule Like-1* (*DSCAML1)*, a gene mutated in patients with autism spectrum disorder, cortical abnormalities, and developmental disorders (Iossifov et al., 2014; Karaca et al., 2015; Deciphering Developmental Disorders Study, 2017). In zebrafish, *dscaml1* is broadly expressed in visual areas and required for visual and visuomotor behaviors, suggesting an underlying visual circuit deficit (Ma et al., 2019). Therefore, we compared the retinofugal connectivity patterns between 5.5 dpf wild-type fish and their *dscaml1* mutant (*dscaml1−/−*) siblings.

We first asked whether loss of *dscaml1* affected the topographical distribution of retinorecipient neurons (340 and 267 in wild type and *dscaml1−/−*, respectively) (Fig. 5). As the initial site of viral infection was in the temporal retina, these retinorecipient neurons likely respond to frontal visual stimulus. The overall distribution was similar between cohorts along the three cardinal axes, but the proportion of retinorecipient cells was significantly shifted in the rostral-caudal and lateral-medial axes. Along the rostral-caudal axis, both excitatory and inhibitory retinorecipient neurons from the *dscaml1*−/− cohort were more rostrally distributed, compared to wild type (p<0.001 and p<0.05 for excitatory and inhibitory neurons, respectively. K-S test). Along the dorsal-ventral axis, the effect of *dscaml1* deficiency was milder. *dscaml1−/−* retinorecipient cells were more dorsally distributed, compared to wild type, but only for excitatory neurons (p<0.05, K-S test). There were no differences in lateral-medial distribution. These results suggest that loss of *dscaml1* may affect the topographic mapping of visual inputs, particularly along the rostral-caudal axes.

**Figure 5.**
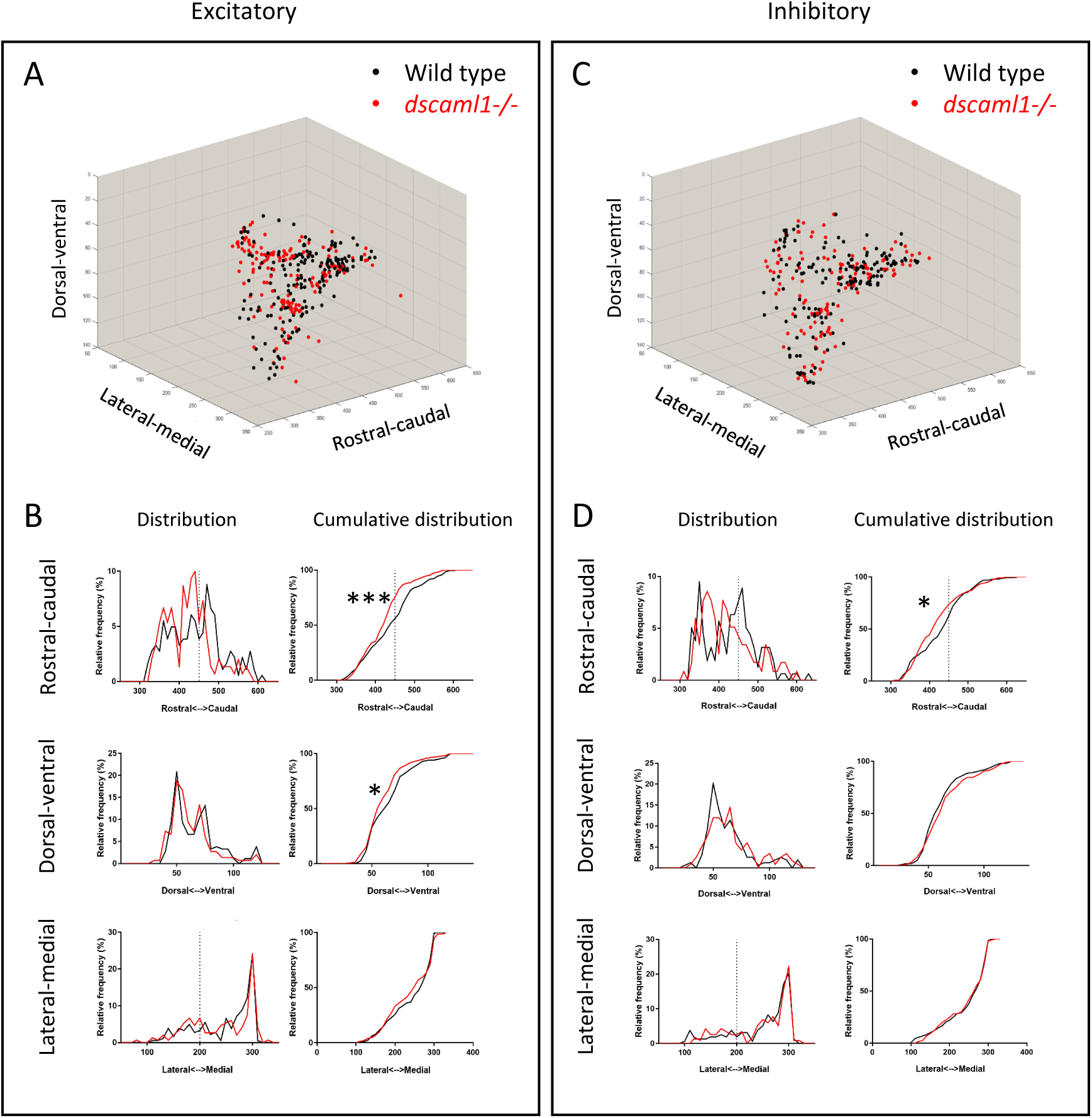
Topographical organization of retinorecipient neurons in wild-type and *dscaml1−/−* cohorts. Graphs show the topographical distribution of excitatory (A, B) and inhibitory neurons (C, D) in wild-type (black, n=24) and *dscaml1−/−* (red, n=26) cohorts. Axes are relative distances (pixels) within the Z-Brain reference brain stack. **A, C,** Three-dimensional distribution of excitatory (A) and inhibitory (C) neurons. **B, D,** Frequency distributions of excitatory (B) and inhibitory (D) neurons in the rostral-caudal, dorsal-ventral, and lateral-medial axes. Dashed lines indicate the boundary between the diencephalon and the mesencephalon. K-S test, *: P<0.05; ***: P<0.001.

Next, we focused on the distribution of retinorecipient cells within specific annotated brain regions. We adapted the Z-Brain quantification tools to measure the sum pixel intensity derived from TRAS-labeled cells for each region (see methods). Among regions defined by anatomy (i.e., not defined by transgene expression), 16 were found to contain, on average, at least 1 retinorecipient cell per animal in the wild-type cohort (Fig. 6A, Supplementary Figure S4). Two major brain divisions, the mesencephalon and rhombencephalon, encompassed all of the retinorecipient cells. Within these divisions, the retinorecipient cells are located within subregions corresponding to known to receive retinofugal input, including the preoptic area, hypothalamus, thalamus, eminentia thalami, pretectum, and optic tectum (tectum neuropil, tectum stratum periventriculare, and medial tectal band) (Burrill and Easter Jr, 1994; Zhang et al., 2017; Helmbrecht et al., 2018; Kramer et al., 2019). We saw no cellular labeling in the olfactory bulb (which innervates the retina), indicating that lentivirus complementation did not facilitate retrograde spread (Li and Dowling, 2000). We also identified several retinorecipient areas that, to the best of our knowledge, had not previously been identified (torus semicircularis, tegmentum, posterior tuberculum).

**Figure 6.**
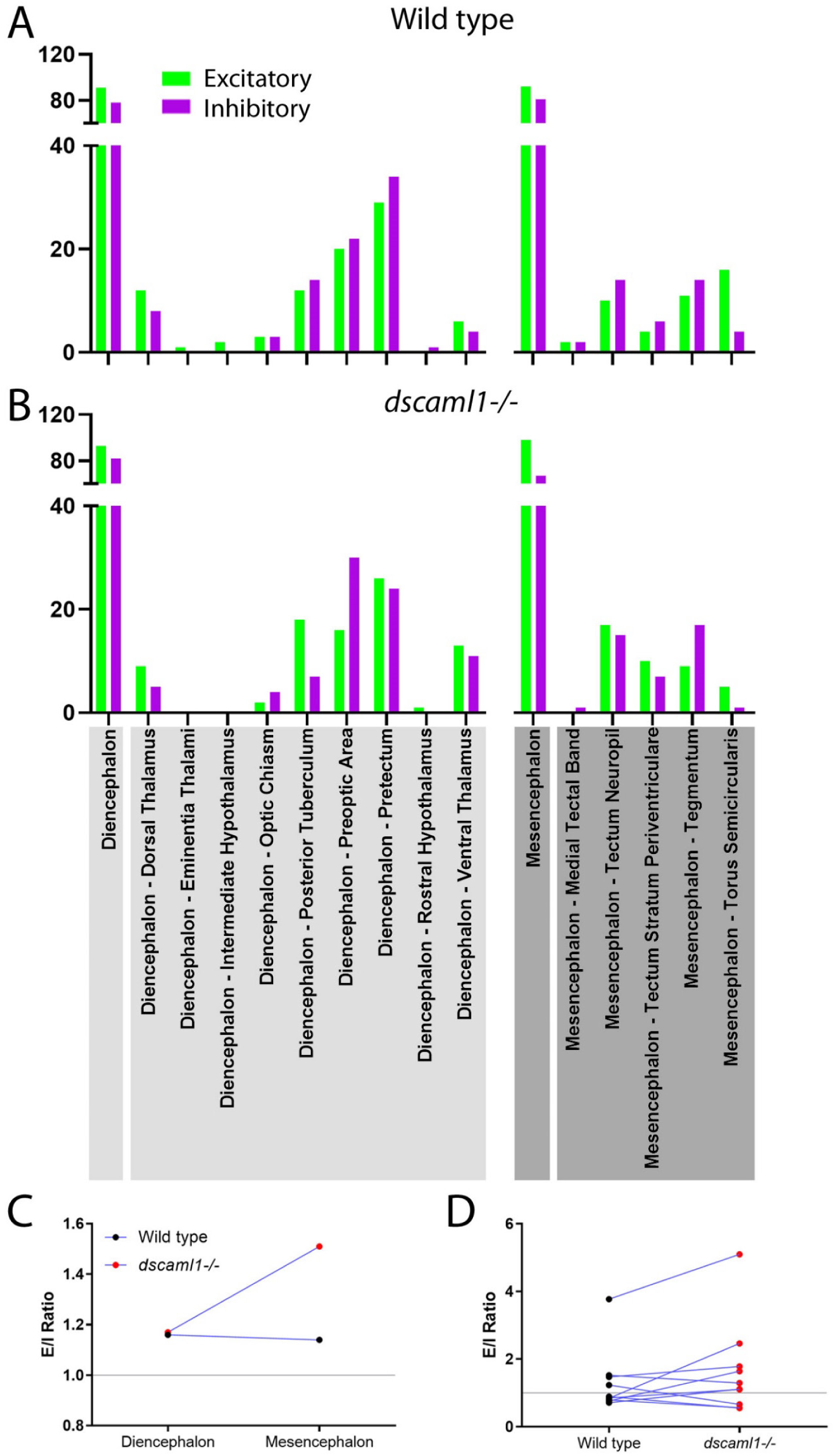
Retinorecipient cell distribution and excitatory-inhibitory balance in major anatomical regions. **A-B,** Axes show estimated cell numbers (converted from intensity signals) in each region in wild-type (A, n=24) and *dscaml1−/−* (B, n=26) cohorts. **C-D,** Excitatory/inhibitory ratio for all labeled non-zero regions in both wild-type (n=24) and *dscaml1−/−* (n=26), listed in **A** and **B**. **C,** Excitatory/inhibitory ratio comparison of diencephalon and mesencephalon. **D,** Excitatory/inhibitory ratio comparison of subregions within diencephalon and mesencephalon.

In general, the same areas were innervated in both wild-type and *dscaml1−/−* cohorts, except for two smaller areas that were not innervated in the *dscaml1−/−* cohort (eminentia thalami and intermediate hypothalamus) (Fig. 6A-B). This result indicates that major retinorecipient areas are innervated by the optic nerve in the *dscaml1−/−* animals. Interestingly, the ratio of excitatory versus inhibitory retinorecipient cells (E/I ratio) was more variable in the *dscaml1−/−* cohort, compared to wild type. This was true both for major brain divisions (diencephalon vs. mesencephalon) and subregions (Fig. 6C-D). A possible explanation for this might be that loss of *dscaml1* affects the targeting of specific cell types within each retinorecipient region.

### TRAS mapping of habenular-recipient neurons

Finally, to test whether TRAS can be applied more generally to other CNS neuronal populations besides RGCs, we examined efferent targeting from the left habenula (Bianco and Wilson, 2009; Amo et al., 2010; Lee et al., 2010; Dreosti et al., 2014; Duboue et al., 2017; Zhang et al., 2017). The bilaterally asymmetrical habenula receives many different sensory cues and is involved in processing social cues, fear learning, and avoidance. The left habenula is known projects to the interpeduncular nucleus (IPN) and superior raphe, providing a suitable pathway to test TRAS mapping (Bianco and Wilson, 2009; Amo et al., 2010). TRAS labeling and image registration were performed as described above, except that virus were injected in the left habenula of wild-type fish. Registered image stacks from five animals with selective left habenula labeling were combined (Fig. 7A-B). We observed consistent labeling of the habenular axon tract and the characteristic annular axon terminals in the IPN. Cell bodies near IPN axon terminals were manually marked (Fig. 7C-D). Consistent with previous findings, habenular target cells we labeled within the IPN and raphe nucleus (Amo et al., 2010). A group of labeled cells is located immediately dorsal to the annotated superior raphe nucleus in Z-Brain, which are likely glial cells. These results demonstrate the general applicability of TRAS for mapping targets of efferent axons.

**Figure 7.**
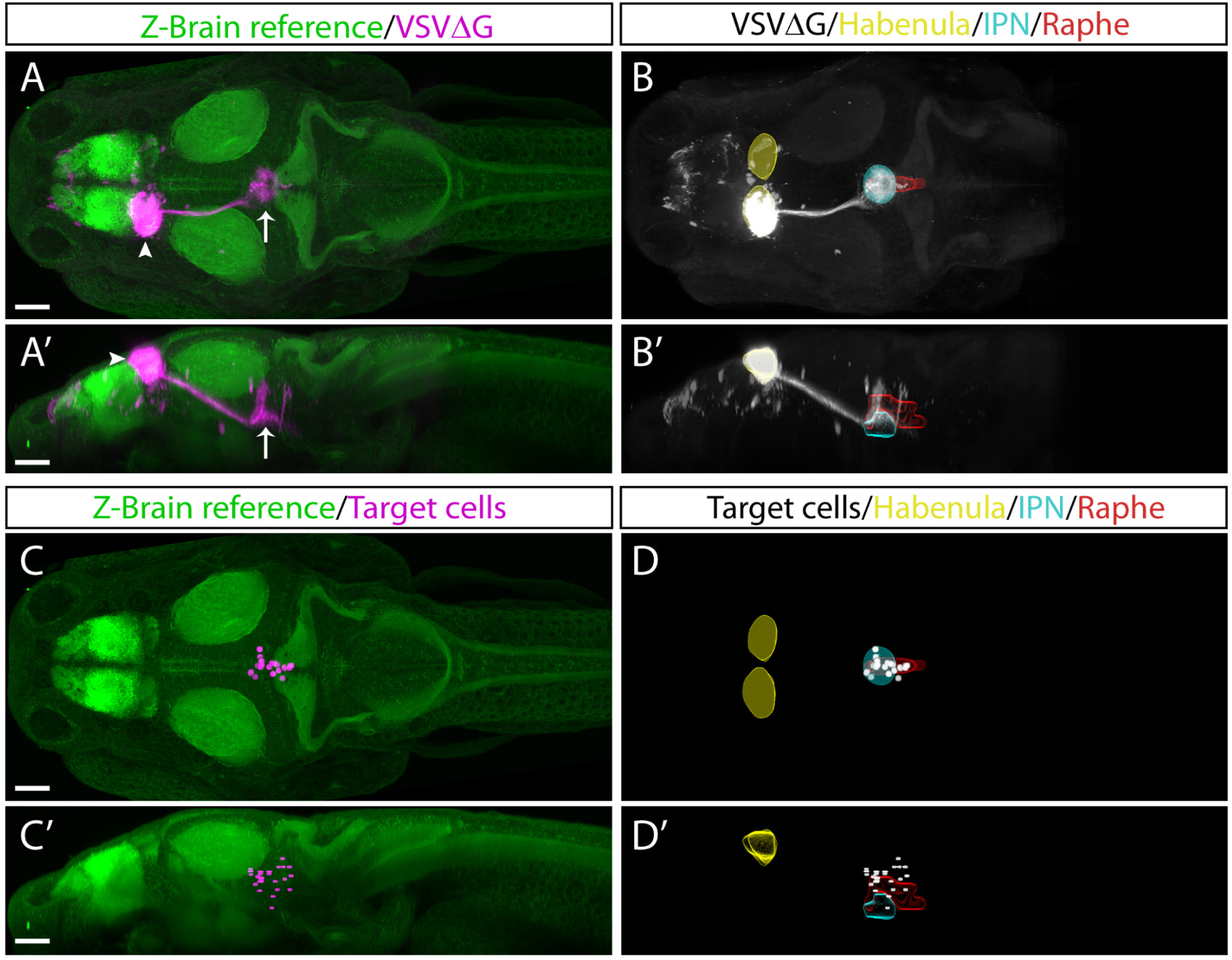
TRAS labeling of habenular target cells. Dorsal views (A-B, C-D) and lateral views (A’-B’, C’-D’) are shown. **A-B,** Maximal projection of registered image stacks from animals with initial infection in the left habenula (arrowhead). Habenular efferent tract projects into the IPN (arrow). RFP expression from VSVΔG infection is shown in magenta (A-A’) or white (B-B’). For anatomical reference, images are overlaid on top of the Z-Brain ERK1/2 reference stack (green, A-A’), or region outlines for the habenula (yellow), IPN (cyan), and raphe nucleus (red) (B-B’). **C-D,** Manually marked habenular target cells (magenta in C-C’, white in D-D’) are overlaid on top of anatomical references, as described for panels A-B. Scale bars are 100 μm. All images are shown at the same scale.

## Discussion

In this study, we developed TRAS, a new method for monosynaptic anterograde labeling in larval zebrafish. This method was applied to the retinofugal pathway and also validated in the habenula efferent pathway. We showed that TRAS could be combined with the Z-Brain image registration and quantitation pipeline to identify changes in retinofugal connectivity patterns caused by the loss of *dscaml1*. These results demonstrate the broad utility of TRAS for neural circuit studies in zebrafish.

### Trans-complementation of VSVΔG by lentivirus

The structure and function of VSV-G have been extensively studied in the context of viral entry, membrane fusion, toxicity, and subcellular transport (Dotti and Simons, 1990; Thomas et al., 1993; Ang et al., 2004; Hoffmann et al., 2010; Albertini et al., 2012; Fossati et al., 2014; Kim et al., 2017). VSV-G also determines the infectivity of VSV-G coated viruses (VSV, RABV, lentivirus, retrovirus), which is crucial for their research and clinical applications (Wickersham et al., 2013; Amirache et al., 2014; Mundell et al., 2015; Kobayashi et al., 2016). Our findings revealed a new aspect of VSV-G function, where VSV-G protein from a different viral species can be recycled to generate infectious VSV.

The spread of VSVΔG from the retina to CNS neurons indicates that VSV-G on the lentivirus surface remained functional after lentivirus infection, and a portion of it was transported anterogradely from the cell body to the axon terminal. At the axon terminal, lentivirus-derived VSV-G was able to re-encapsulate the VSV nucleocapsid and mediate subsequent infection. The strategy of using lentivirus as a tool for glycoprotein complementation could potentially be applied more broadly. For example, current strategies for RABV monosynaptic tracing utilizes AAV to express RABV glycoprotein, which usually takes several weeks for sufficient glycoprotein expression (Miyamichi et al., 2011). It will be interesting to test whether rabies glycoprotein-coated lentivirus could be a more expedient method to provide glycoprotein for retrograde tracing.

### Applying TRAS for zebrafish neural connectivity analysis

Advances in viral engineering have led to new neural circuit tracing strategies utilizing replication-incompetent viruses (e.g., RABV, AAV, HSV) that are safer to use, less toxic to host cells, and have restricted (mostly monosynaptic) spread (Wickersham et al., 2007; Zingg et al., 2017; Chatterjee et al., 2018; Beier, 2019). Unfortunately, many of the transsynaptic viruses used in mammalian systems either do not infect zebrafish (e.g., AAV) or have low efficiency for transsynaptic spread (e.g., RABV) (Zhu et al., 2009; Dohaku et al., 2019). VSV, in contrast, can infect larval zebrafish and spreads robustly both anterogradely and retrogradely. However, replication-competent VSV has high cytotoxicity and can spread across multiple synapses, making it difficult to distinguish between direct versus indirect connections (Mundell et al., 2015).

To address these limitations and provide a tool for neural circuit mapping for larval zebrafish, we developed TRAS. TRAS utilizes recombinant VSV with genomic deletion of the glycoprotein gene (VSVΔG). VSVΔG can infect cells at the injection site but cannot spread. Although wild-type VSV does not cause serious illness to humans, the use of VSVΔG further reduces the risk of exposure (Spickler, 2016). The lack of VSV-G expression from the viral genome also helps reduces toxicity to the host cell, as long-term VSV-G expression is known to be cytotoxic (Yee et al., 1994). To complement VSVΔG, we directly provided VSV-G protein, utilizing lentivirus as the transducing reagent. Compared to transgenic or virus-induced expression, this approach is rapid and transient, therefore minimizing the cellular exposure to VSV-G. Both VSVΔG and VSV-G coated lentivirus are available from a commercial source, making TRAS an easy method to adopt in a typical neuroscience laboratory.

To extend the utility of TRAS, we developed procedures to register brain images to the Z-Brain anatomical template (Randlett et al., 2015). The combination of neural circuit tracing within a standard 3D-brain atlas is the current state of the art approach for understanding neural network connections, both in zebrafish and mammalian models (Watabe-Uchida et al., 2012; Oh et al., 2014; Helmbrecht et al., 2018; Kramer et al., 2019; Kunst et al., 2019). This approach provides a more objective way to map cells and pathways onto specific brain regions across different experimental animals and promotes cross-referencing between research findings.

We demonstrated that TRAS and Z-Brain could be used for neural circuit mapping in efferent pathways originating from the retina and the left habenula. These are two of the better-studied pathways in larval zebrafish, which allowed us to assess the specificity of TRAS for anterograde labeling of direct postsynaptic targets. Overall, TRAS identified all of the target regions described in previous studies, which gives confidence to the future application of TRAS to map unknown neural connections in zebrafish. Furthermore, given that VSV is also an anterograde tracer in mice and chicken, it will be interesting to test whether TRAS can be applied to these experimental systems for neural circuit mapping (Mundell et al., 2015).

### Limitations and the future development of TRAS

While TRAS offers many advantages as a neural circuit mapping tool, it is important to note some of the limitations of the technique. These are also areas with potential for further technological development. First, since VSV-G binds to a receptor that is widely expressed (LDL receptor) (Finkelshtein et al., 2013), VSV-G coated viruses (e.g., VSVΔG and lentivirus) can infect most cell types. Therefore, the specificity of TRAS depends on precise injection into the brain region of interest. For brain regions smaller than the habenula, a compound microscope with DIC optics would be necessary. To restrict infection to a particular cell type, it may be possible to make use of ASLV-A pseudotyped VSVΔG that can selectively target neurons expressing an exogenous receptor, TVA (Beier et al., 2011b; Dohaku et al., 2019). However, potential interactions between virions with different envelope glycoproteins may interfere with the specificity of VSVΔG infection (Beier et al., 2011a)

Second, while VSVΔG by itself cannot spread after initial infection, it can still replicate and change the metabolism of the host cell. For instance, the VSV M protein is capable of altering host cell transcription and translation. Chronic VSVΔG infection would likely affect the survival of infected neurons and impair its neurophysiological functions. Several approaches for reducing the toxicity of RABV have been reported recently to reduce the function or expression of viral proteins, such as destabilizing the RABV nucleoprotein or deleting the RABV L gene (Ciabatti et al., 2017; Chatterjee et al., 2018). Similar manipulations may also reduce the toxicity of VSV.

Third, we observed TRAS labeling of cells that do not express the *HuC:H2B-GCaMP6f* transgene (non-GCaMP cells), which may be glial cells. While not frequently discussed, viral transmission from neuron to glia does occur for most viral transsynaptic tracers, including PRBV, HSV, RABV, and VSV (Beier, 2019). In a replication-competent virus, the infection of glia and subsequent spread to neurons would make it challenging to infer connectivity. This issue is circumvented in TRAS, as VSVΔG cannot spread after glial infection (due to the lack of VSV-G). Nevertheless, this highlights the importance of distinguishing bona fide neurons versus other cell types in any type of transsynaptic labeling study.

Lastly, quantitation by Z-Brain depends on morphing and registration of image stacks to a reference template, followed by manual identification of labeled neurons. This approach is suitable to test the effects of single genes or pathological states, but likely too laborious as a screening tool to identify candidate genes or screen drugs. Selective fluorescent labeling of neuronal cell bodies (without labeling neurites) and automation of cell detection would be a crucial next step to improve the utility of TRAS.

### Connectivity patterns associated with *dscaml1* deficiency

The ability to quantitate efferent connections prompted us to investigate whether TRAS can be used to identify connectivity deficits caused by *dscaml1* deficiency. As mentioned previously, human DSCAML1 mutations are believed to be causative for neurodevelopmental disorders. Additionally, our recent work has found that loss of *dscaml1* significantly impaired visuomotor function associated with light perception and eye movements, suggesting a possible underlying deficit along the visual pathway (Ma et al., 2019).

Using TRAS and Z-Brain quantification, we found that *dscaml1* deficiency might have a role in refining the retinofugal topography and cell-type specificity. On a broader scale, we saw similar patterns of topographic and region-specific projections between wild-type and *dscaml1−/−* cohorts (Fig. 6, 7). This indicated that RGC axonal targeting was mostly intact in the *dscaml1* mutants. Interestingly, there was a significant rostral shift in the position in both excitatory and inhibitory retinorecipient cells. Given that RGC axon terminals and retinorecipient cells are both topographically organized, this shift in positioning may result in diminished spatial perception (Stuermer, 1988; Muto et al., 2013; Robles et al., 2014). We also observed a trend for altered ratio of excitatory versus inhibitory retinorecipient cells in the *dscaml1* mutants. For example, the preoptic area and tegementum have higher a ratio of inhibitory neurons in the *dscaml1* mutants, compared to wild type. This putative change in retinofugal target selection could lead to network changes and altered visual response. Further physiological studies will be needed to formally test whether *dscaml1* affects spatial perception and the excitatory/inhibitory balance in the visual pathway.

### Conclusions

Here we present the development of a new technique (TRAS) that is suitable for mapping neural connectivity in zebrafish. TRAS makes use of a novel lentivirus trans-complementation approach to enable restricted anterograde transneuronal spread by recombinant VSV. We have validated this method in two efferent pathways and identified potential connectivity pattern changes caused by a genetic deficiency in *dscaml1*, a neuronal cell adhesion molecule associated with human neurodevelopmental disorders. The ability of TRAS to map structural connectivity would enable the discovery of new neural connections and complement existing brain mapping efforts.

## Acknowledgments

This work was supported by funding from the National Institutes of Health (R01 EY024844 to Y.A.P.), Augusta University, and Virginia Tech. We thank the animal care staff at Augusta University and Virginia Tech for animal husbandry, Owen Randlett for help with Z-Brain analysis, Didem Gotz Ayturk, Xiang Ma, and Constance Cepko for reagents and technical expertise for virus preparation, and members of the Pan lab for helpful suggestions on the manuscript.

## Author Contributions

M.M., S.K., and Y.A.P. conceived the study, performed the experiments, and analyzed the data. S.K. prepared and characterized recombinant viruses. M.M. and Y.A.P. wrote the manuscript, with contributions from S.K.

## Videos

**Supplementary Video 1.** Image stack of TRAS-labeled zebrafish larva, three days after the initial infection. VSVΔG labeling is shown in magenta and *vglut2a:GFP* labeling in green.

**Supplementary Video 2.** Excitatory (green) and inhibitory (magenta) retinorecipient cells in wild type (left) and *dscaml1−/−* (right) cohorts. ERK1/2 immunolabeling (white) from Randlett et al. 2015 is overlaid to serve as an anatomical reference.

## Data Availability

The image stacks and other data supporting the findings of this study are available from the corresponding author upon reasonable request.

## Code Availability

Custom Fiji and MATLAB scripts used of this study are available from the corresponding author upon reasonable request.

---

**Supplementary Figure S1.**
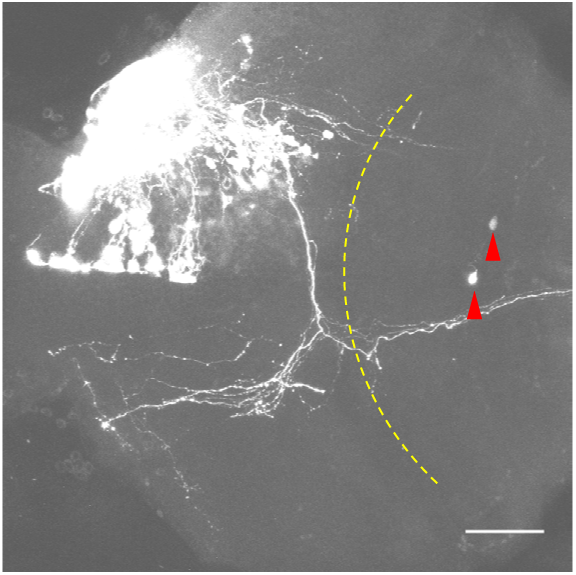
Examples of secondary TRAS labeling. Red arrowheads indicate secondary spread due to a higher titer of virus. Yellow dash line marks the border of mesencephalon and cerebellum. Scale bar: 50 μm.

**Supplementary Figure S2.**
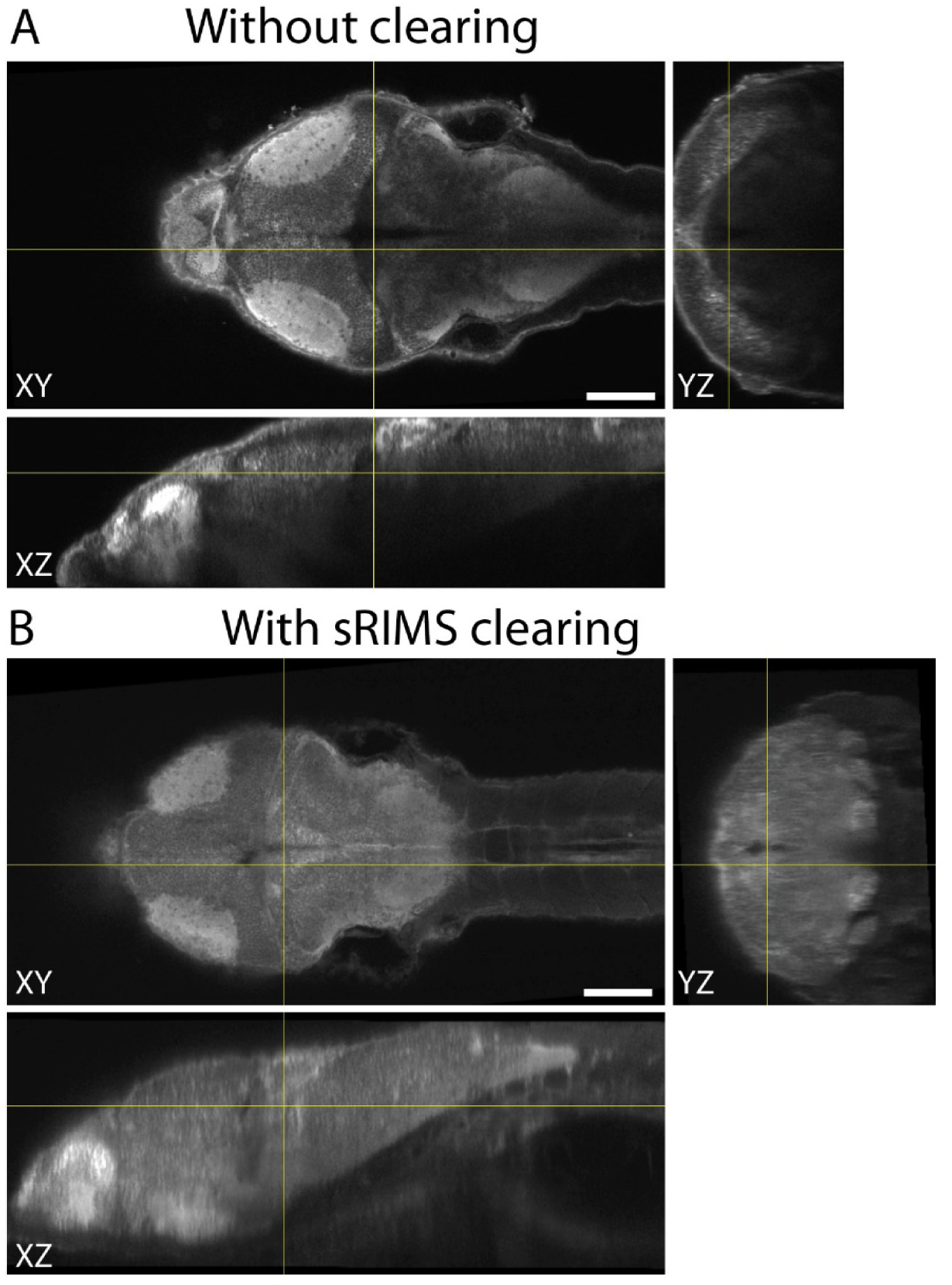
Tissue clearing with sRIMS solution. Whole-mount ERK1/2 immunolabeling without (A) or with sRIMS clearing (B). Orthogonal views (XY, XZ, and YZ) of confocal image stacks are shown, centered just bellowed the cerebellum (intersect of yellow lines). Ventral structures are not visible without sRIMS clearing. Scale bars are 100 μm.

**Supplementary Figure S3.**
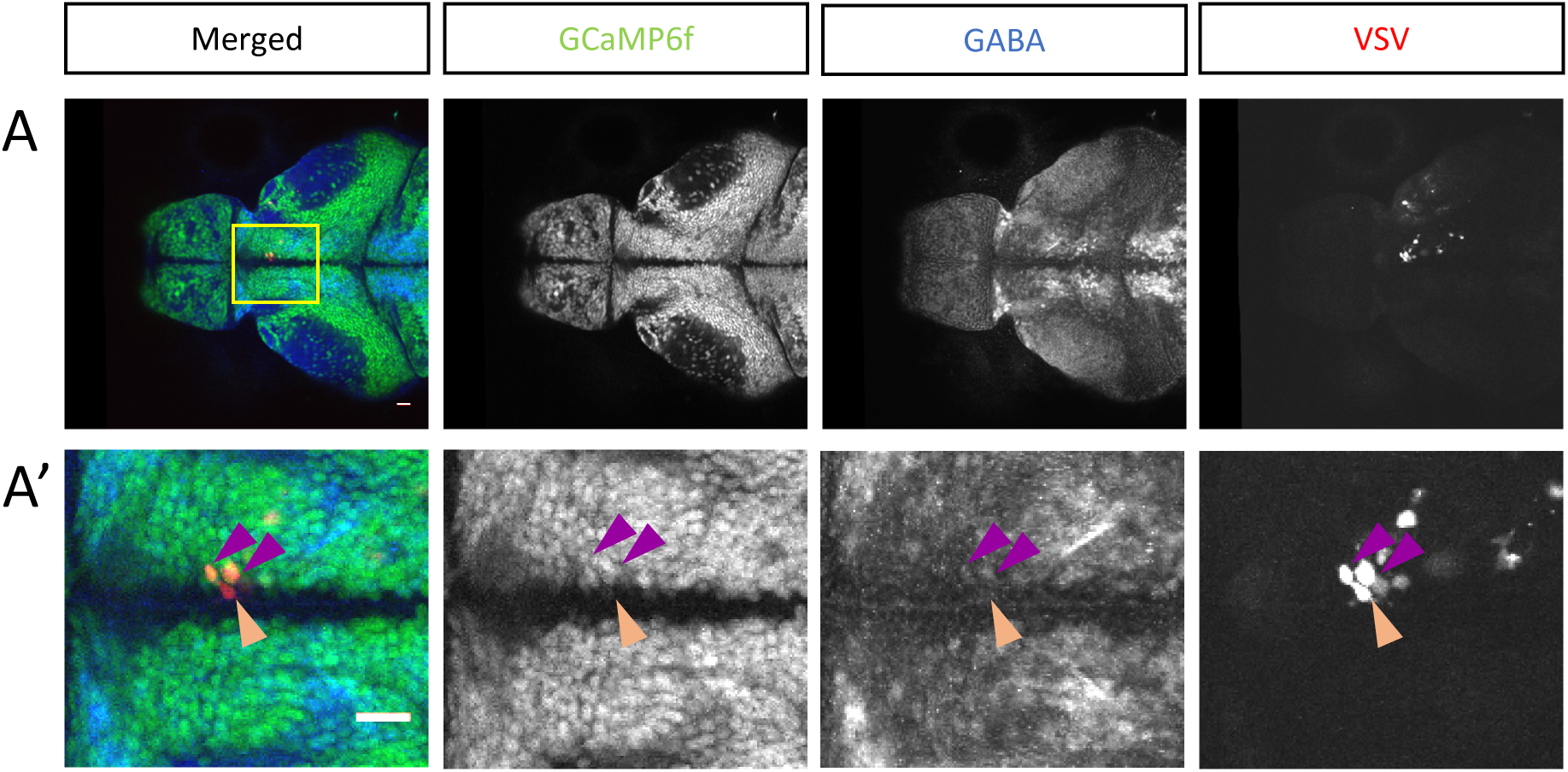
TRAS labeling of non-neuronal cells. A single confocal imaging plane is shown, with merged, GCaMPf, GABA, and VSVΔG channels as indicated. Boxed area in A is shown in higher magnification in A’. The purple arrowheads mark two GCaMP6f+/GABA+/VSVΔG+ inhibitory neurons. The orange arrowhead marks a GCaMP6f-/GABA-/VSVΔG+ cell. Scale bars are 100 μm.

**Supplementary Figure S4.**
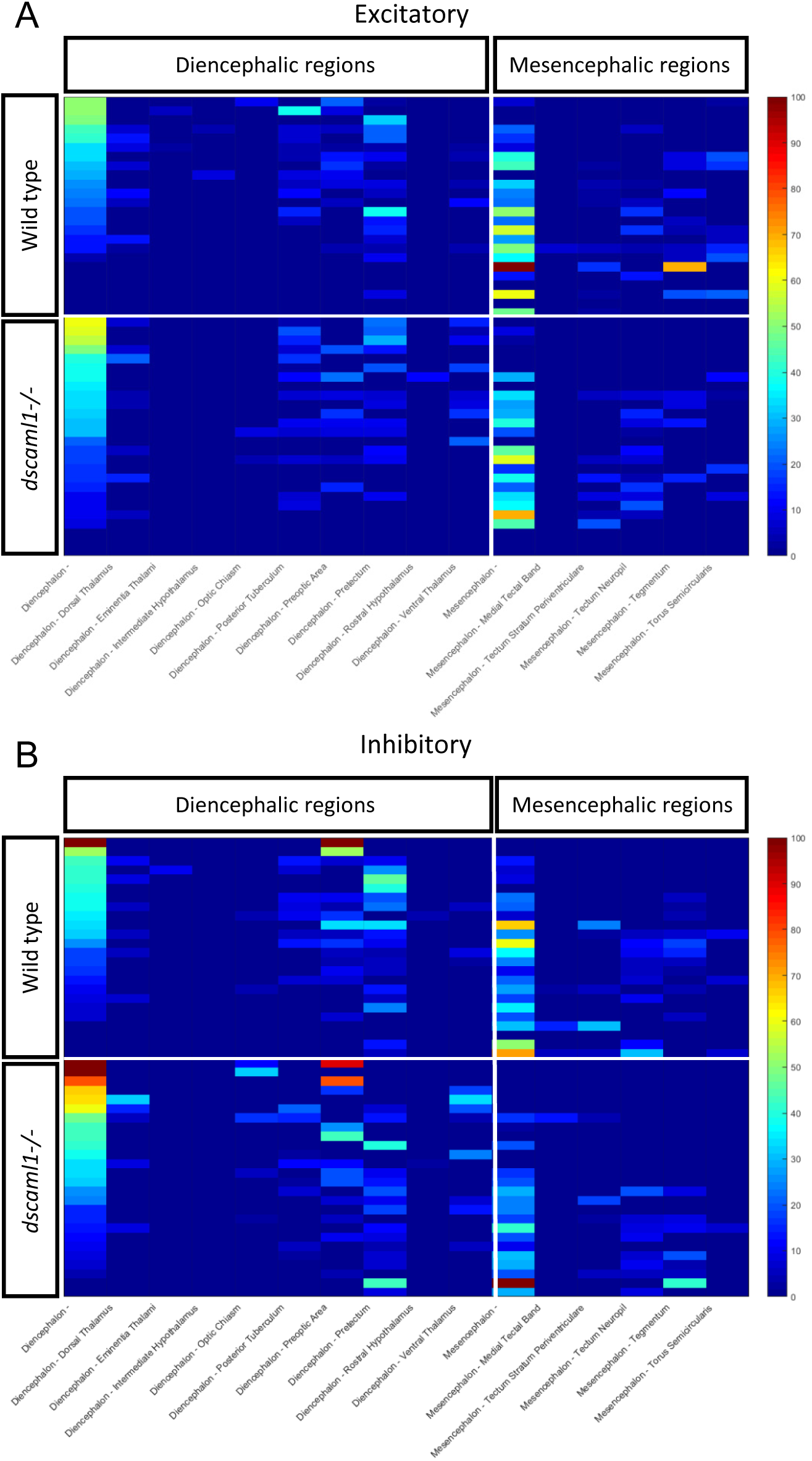
Signals detected via Z-brain in wild-type and *dscaml1−/−* fish. **A-B,** Heat map of normalized signals in major anatomical regions from Z-brain showing normalized signals detected within each sample used for analysis, wild-type (n=24) and *dscaml1−/−* (n=26), and their corresponding neuron types **A** is excitatory while **B** is inhibitory. All signals were descended aligning to Diencephalon.

